# Laboratory adaptation and complete genome assembly of a Beposo, Ghana strain of the human hookworm *Necator americanus*

**DOI:** 10.64898/2026.06.10.728644

**Authors:** Lisa M. Harrison, Kaylee S. Herzog, Dickson Osabutey, Mavis Konoma, Emma Allen, Kelly Hagadorn, Santosh George, Richard Bungiro, Claudia Gaither, Carol Mariani, Margaret K. Corley, Adalgisa Caccone, Michael Wilson, Joseph R. Fauver, Michael Cappello

## Abstract

Laboratory models are invaluable tools for studying parasite biology and pathogenesis, especially for helminth infections. However, the complex life cycles and frequently narrow host specificity of helminths present challenges to maintaining access to critical parasite material in a laboratory setting. This is especially true of *Necator americanus*, the most common species of hookworm that infects humans globally. Here we report the successful laboratory adaptation of an African strain of *N. americanus*, originally isolated from infected individuals in Beposo, Ghana. The Beposo strain has been successfully passaged across 9 generations in Golden Syrian hamsters maintained on oral dexamethasone. Differential susceptibility to mebendazole and albendazole was evaluated using an egg hatch assay, and DNA sequencing of the beta-tubulin isotype 1 gene did not identify known resistance-associated mutations in the endemic strain. Sequencing of the mitochondrial *COX1* gene revealed that specimens of *N. americanus* from Ghana, along with reported sequences from Togo, are distinct from those from South America and Asia. Complementary microsatellite-based population analysis revealed substantial genetic variation in the founding parasite population. To further characterize the novel Beposo strain, a draft hybrid genome assembly was generated from genomic DNA extracted from a single adult male worm via an optimized Oxford Nanopore Technologies MinION library preparation approach tailored to low-input sample types. This high-quality assembly, including a complete mitogenome, is 202.8Mb in 950 contigs with an N50 >449 kb. It contains >95% of conserved nematode orthologs in complete single copy and is estimated by homology-based gene prediction to contain 12,804 genes. This study represents the first comprehensive characterization of a strain of *N. americanus* originating in Africa that has been successfully adapted to a laboratory animal model.

## Introduction

According to the recent estimates from the World Health Organization [1], more than 1 billion people are infected with soil-transmitted helminths (STHs), with the highest prevalence among those who live in rural areas and lack adequate access to sanitation or clean water. Despite longstanding recognition of the global health significance of STH infections, including decades of commitment to reducing the burden of disease among the most vulnerable populations, parasitic worms remain significant causes of malnutrition and anemia in low and middle income countries [2–6].

Among the STHs, hookworms infect approximately 500 million people worldwide and are a significant cause of anemia, growth delay and poor birth outcomes in vulnerable populations, including children and women of child-bearing age [7–10]. Infection typically occurs through skin contact with infectious third stage larvae (L3) in soil contaminated with human feces. Once within the skin, hookworms invade small veins or lymphatic vessels and are carried with the circulation to the heart and lungs. After lodging in alveolar capillaries, the migrating larvae cross from the blood vessels into the small airways, eventually crawling up the respiratory tree where they are swallowed and carried by peristalsis to the small intestine. After completing their final molt to the adult stage, hookworms attach to the intestinal mucosa and feed on blood from lacerated blood vessels, a process facilitated by secretion of anticoagulants, anti-inflammatory molecules and proteases.

A major impediment to the study of hookworms and other STHs is the limited availability of sustainable animal models to characterize parasite biology, disease pathogenesis and host-parasite interactions, which has slowed development of drugs, diagnostics and vaccines needed to treat, monitor and prevent hookworm infection in endemic populations [11–13]. Like many parasites, helminths typically have narrow host species specificity, limiting the number of animal models available to researchers. For example, mice and rats are not fully permissive hosts for the most common hookworm species that infect humans (*Necator americanus*, *Ancylostoma duodenale*), thereby limiting the number of immunologic reagents and genetic tools that can be applied to studying these parasites *in vivo*. The most reliable and cost-effective host species for hookworm research has proven to be the outbred hamster (*Mesocricetus auratus*), which has been adapted as a fully permissive host for *N. americanus* and the zoonotic hookworm, *A. ceylanicum*, which is an emerging parasite of humans in Asia [14–19].

While the hamster model represents a viable alternative to mice and rats, to date there have been very few reports of successful adaptation of a human field isolate and maintenance of the life cycle in a laboratory setting [14, 20–22]. Moreover, we found no reports describing the sustained passage of a human hookworm strain originating in Africa, where the burden of disease remains a significant public health challenge. We report here the successful laboratory propagation and genomic characterization of an endemic human strain of *N. americanus* originally isolated in Beposo, Ghana. We also analyze the phylogenetic position of the Ghana strains in relation to geographically distinct isolates using mitochondrial *COX1* gene sequences, assess levels of nuclear variation of the Ghana strains using 21 microsatellite loci, and evaluated *in vitro* their susceptibility to benzimidazole anthelminthics (albendazole, mebendazole). Finally, using a single adult worm, we have assembled the first complete hookworm genome from West Africa, providing a valuable resource for future genomic studies of this globally important human parasite.

## Methods

### Ethics statement

Informed consent was obtained from all human study subjects enrolled in the field survey conducted in Beposo, Ghana. Prior to enrolling human subjects, research study protocols were independently reviewed and approved by the IRB committees of Yale University and the Noguchi Memorial Institute for Medical Research at University of Ghana. Animal husbandry was provided by the Yale Animal Resource Center and animal experimentation protocols approved by the Yale Animal Care and Use Committee prior to study initiation.

### Laboratory passage of Beposo field isolate of *Necator americanus*

From August 2-4, 2019, a targeted field survey was conducted in order to isolate hookworm larvae from human subjects (n=300) living in Beposo, which is located in the Pru West District in the Bono-East region of Ghana. Fecal samples from study subjects found to be hookworm infected using Kato-Katz microscopy were combined, mixed with bone char coal (Ebonex, Melvindale, Michigan) and cultured at ambient temperature for 10-14 days. Viable L3 were collected and concentrated using a Baermann funnel apparatus [23, 24]. Species-specific PCR of cultured larvae confirmed monocultures of *N. americanus* [25, 26].

*In vivo* laboratory propagation of the Beposo *N. americanus* strain of hookworm was achieved through serial infection of weanling (21 days old) male Golden Syrian Hamsters (*Mesocricetus auratus*) of the outbred HsdHan: AURA (LVG) strain (Envigo, Indianapolis, IN). Prior to the initial and subsequent infections with *N. americanus* L3, hamsters (n=6 per cage group) were provided drinking water *ad libitum* containing dexamethasone (2.5µg/ml) for 4 days. Animals were then injected subcutaneously with 300 L3 suspended in PBS and monitored for fecal egg excretion using McMaster slide microscopy [27]. Once patent infection was confirmed, hamster feces was cultured and infectious larvae isolated to initiate subsequent passages in weanling hamsters.

### Benzimidazole susceptibility of *Necator americanus* field isolates

Susceptibility of the Beposo strain to benzimidazole anthelminthics was measured and compared to that of *Ancylostoma ceylanicum* using a previously described egg hatch assay [28, 29]. Pooled fecal samples were collected from Golden Syrian hamsters following infection with L3s (*A. ceylanicum* or *N. americanus*) as previously described [29, 30]. Separately for each species, hookworm eggs were purified from hamster feces using a density floatation method and resuspended in water. Eggs were pipetted into 96-well plates at a concentration of 100 eggs per well, followed by the addition of benzimidazole compound dissolved in DMSO. Each drug was tested in duplicate at final concentrations ranging from 0–10 μM. Egg hatch assay (EHA) cultures were incubated for 24 hrs at ambient temperature, at which time the numbers of larvae and unhatched eggs were counted by light microscopy and percent egg hatch inhibition values were calculated as:

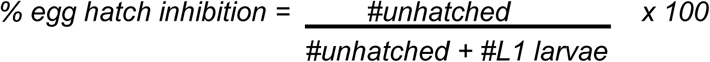

### DNA sequencing of partial beta-tubulin isotype 1

Genomic DNA was extracted from hookworms by incubating individual adult parasites (n=19) in 50µl lysis buffer (50 mM KCl, 10 mM Tris pH 8.3, 2.5 mM MgCl_2_, 0.45% Tween-20, 0.01% Gelatin) and proteinase K (100µg/ml) overnight at 37°C. Proteinase K was inactivated by heating samples for 20 minutes at 70°C [31, 32]. PCR was used to amplify regions containing previously identified benzimidazole resistance mutations (F167Y, E198A, and F200Y) in the beta-tubulin isotype 1 gene from *N. americanus* (GenBank: EF392851.1). Amplicon 1 corresponded to nucleotides 948-1743 of the beta-tubulin gene, while amplicon 2 represented nucleotides 1621-2143. PCR products of the expected size were purified from agarose gel slices using the QIAquick PCR & Gel Cleanup Kit (QIAGEN), and sequencing of purified PCR products was carried out by the Yale Center for Genome Analysis (YCGA).

### DNA sequencing of the mitochondrial cytochrome c oxidase subunit (*COX1*)

Amplification of the *COX1* gene was carried out as previously described [31, 33] to validate the presence of *N. americanus* larvae isolated from hamster fecal cultures. *Necator americanus* genomic DNA (gDNA) previously isolated from human subjects in Kintampo North Municipality, Ghana [34–36] was used as a positive control, along with no template (negative) controls in each assay. PCR products were purified and sequenced by the YCGA as described above.

### Phylogenetic analysis of partial *COX1* sequences

Partial *COX1* sequences were analyzed from adult hookworms removed from hamsters following initial passage of the original field-cultured L3 from Beposo, Ghana (n=21) and the Beposo strain mitogenome assembly (n=1; see Methods below). Additional ingroup sequences for *N. americanus* from other hookworm endemic regions (Togo, Brazil, China; n=20), as well as an outgroup sequence from *Ancylostoma duodenale*, were downloaded from GenBank. Sequences were aligned using MAFFT and manually trimmed in Geneious 2022.2.1. A maximum likelihood topology was generated using IQ-TREE v. 2.3.4 [37, 38]. Model selection was performed by ModelFinder in IQ-TREE [38, 39] and nodal support values were generated through 1,000 ultrafast bootstrap (UFBS) replicates [40].

### Characterization of di-, tri-, and tetranucleotide microsatellite markers of Beposo strain

*In silico* analysis was used to identify microsatellite markers from the previously published genome of *N. americanus* (BioProject PRJNA72135). The program msatcommander (a BioPython program) was used to identify di-, tri- and tetranucleotide microsatellite repeats of 100, 200, 300 and 400 base pair fragment sizes [41, 42]. The entire 245Mb genome sequence (BioProject PRJNA72135) was uploaded in FASTA format to msatcommander revealing 22,632 microsatellite regions. From this initial screen, a subset of 40 microsatellites were shortlisted (**S1 Table**) based on varying repeat sequences and fragment sizes. These loci were targeted for PCR amplification. Primer sequences and characteristics of validated loci are provided in **S2 Table**.

PCR screening was carried out to determine variability between 20 individual F0 adult hookworms using gDNA samples extracted from single worms as previously described [34–36]. The genomic template concentration was quantified using a Qubit 2.0 fluorometer (Life Technologies, USA) prior to PCR amplification. A touchdown PCR was conducted in a 10µL reaction mixture containing the TypeIt Microsatellite PCR Master Mix (Qiagen), 2µM of forward primer (Integrated DNA Technologies) 10µM of reverse primer. Both primers were tagged to allow multiplexing, with fluorescently labelled M13 universal sequence on the 5’ end of the forward primers and with GTTT pigtail on the 5’ end of the reverse primers (**S2 Table**). Thermocycling conditions were the same for all primer sets (95°C for 5 minutes; 12 cycles of 95°C for 30 seconds, 65°C for 30 seconds, 72°C for 30 seconds; 33 cycles of 95°C for 30 seconds, 56°C for 30 seconds, 72°C for 30 seconds; and 1 final extension cycle of 60°C for 30 minutes). Fragment analysis of PCR products was carried out using an Applied Biosystems 3730xl DNA Genetic Analyzer with a GeneScan Liz-500 internal size standard (Applied Biosystems) at the DNA Analysis Facility, Yale University.

Microsatellite peaks were scored manually in GeneMarker v2.4.0 (SoftGenetics), and all allele scores were verified by a second observer to ensure consistency. GeneAlEx 6.5 was used to calculate the number of alleles, as well as the observed (Ho) and expected (He) heterozygosity per locus. GENEPOP V4 was used to perform a Hardy-Weinberg exact test for each locus. MICRO-CHECKER 3.2.2 [43] was used to check for null alleles.

### Sample collection and DNA extraction for genome assembly

A single adult male of *N. americanus* obtained from the first-generation passage through hamsters was preserved in RNAlater (Sigma-Aldrich, USA) and used for genome assembly. Total nucleic acid was extracted using the Quick-DNA HMW MagBead Kit (Zymo Research, USA) according to an optimized protocol as previously described [44]. DNA concentration and fragment length were assessed using the Qubit 1x dsDNA High Sensitivity Assay (Thermofisher Scientific, USA) and the Agilent 2200 TapeStation System for Genomic DNA (Agilent, USA).

### Library preparation & next-generation sequencing for genome assembly

Extracted nucleic acid from the adult male of *N. americanus* was split into two aliquots: one each for Oxford Nanopore Technologies (ONT) MinION and Illumina library preparation and sequencing. A MinION library was prepared with the Ligation Sequencing Kit (ONT, UK) with modifications to preserve nucleic acid quantities through library preparation as previously described for *Brugia malayi* and *Trichuris trichiura* [44]. The final library was loaded onto an ONT MinION R10.4.1 flow cell and sequenced using MinKNOW software v. 22.12.7 (ONT) with pore scans every 1.5 hr. Following ∼30.5 hr of sequencing, remaining library was removed from the flow cell via the SpotON port and cleaned up following the ONT library recovery from flow cells protocol (LIR-9178) with the following modifications: Once removed from the flow cell, the library was bound to AMPure XP beads (Beckman Coulter, USA) for 1 hr on a rotating mixer at RT, washed with Short Fragment Buffer (ONT, UK), eluted at 37°C for 2 hr in a heated dry bath with occasional flick-mixing, and then eluted at RT overnight. The entire volume of eluted library was then re-sequenced on a new R10.4.1 flow cell for ∼5.5 hr using the same run settings described above. An Illumina library was prepared for sequencing at the YCGA using the xGen cfDNA & FFPE DNA Library Prep Kit with unique dual indexing (Integrated DNA Techonologies, USA). The library was sequenced on an Illumina NovaSeq 6000 using 2×150 cycles targeting 50× coverage of the *N. americanus* genome.

### Genome and mitogenome assembly

Methods for quality control of MinION and Illumina read data, estimation of genome size and heterozygosity, and hybrid genome assembly with MaSuRCA v 4.1.0 followed Herzog et al. [44]. To assemble the mitogenome, quality-controlled MinION and Illumina reads were separately mapped to the reference mitogenome for *N. americanus* from Togo (GenBank No. AJ556134) [45] using Minimap2 v. 2.16 [46] and BWA v. 0.7.17 [47] respectively. SAMtools v. 1.9 [48] was then used to generate sorted BAM files from each mapping. Consensus sequences were generated from each BAM file using Geneious Prime v. 2022.0.1 aligned to one another, and manually compared to produce a final hybrid mitogenome sequence. The mitogenome was annotated using the “Annotate from” function in Geneious with the annotated mitogenome from China (GenBank No. AJ417719) [49] as the reference.

### Assessment of genome contiguity, completeness, and gene content

Genome contiguity was assessed with QUAST v 5.0.2 [50]. Genome completeness was assessed with miniBUSCO v. 0.2. using the Nematoda ortholog database (nematodoa_odb10) [51]. Homology-based gene prediction was performed using GeMoMA v 1.9 [52] with the assembly and annotation of Tang et al [53] as a reference. The resulting GFF file was summarized using AGAT v 0.7.0 [54].

## Results

### Sequential passage of the Beposo strain of *Necator americanus* in hamsters

A total of 300 study subjects living in Beposo, Ghana were recruited over 3 consecutive days in August 2019 for a cross-sectional survey of hookworm infection. Up to 10 subjects were enrolled from each of 30 households, with a goal of obtaining a representative sample from the community. After providing informed consent, each study subject submitted a fresh fecal sample, which was evaluated for the presence of hookworm eggs using Kato-Katz microscopy. Baermann cultures prepared from pooled Kato-Katz positive fecal samples produced approximately 5,000 viable L3, which were transported to New Haven, CT USA. Aliquots of L3 were confirmed to be *N. americanus* by *COX1* sequencing.

Fecal egg excretion in hamsters infected with 300 *N. americanus* L3 was monitored on a regular basis across the first 9 passages (F0-F8) in the hamster model. As shown in **Fig 1**, hamsters infected with *N. americanus* L3 transported from the field yielded a patent infection that was first detected by fecal microscopy at day 51 days post-infection (DPI). Peak egg excretion was observed at 56 DPI, with no further egg excretion detected after 62 DPI. Subsequent passages were marked by incremental increases in both the peak and duration of fecal egg excretion. For example, from F0 to F4 passages, peak egg excretion increased from 160 eggs per gram (EPG) to 1007 EPG in hamsters receiving 2.5 µg/ml of dexamethasone in drinking water.

**Fig 1.**
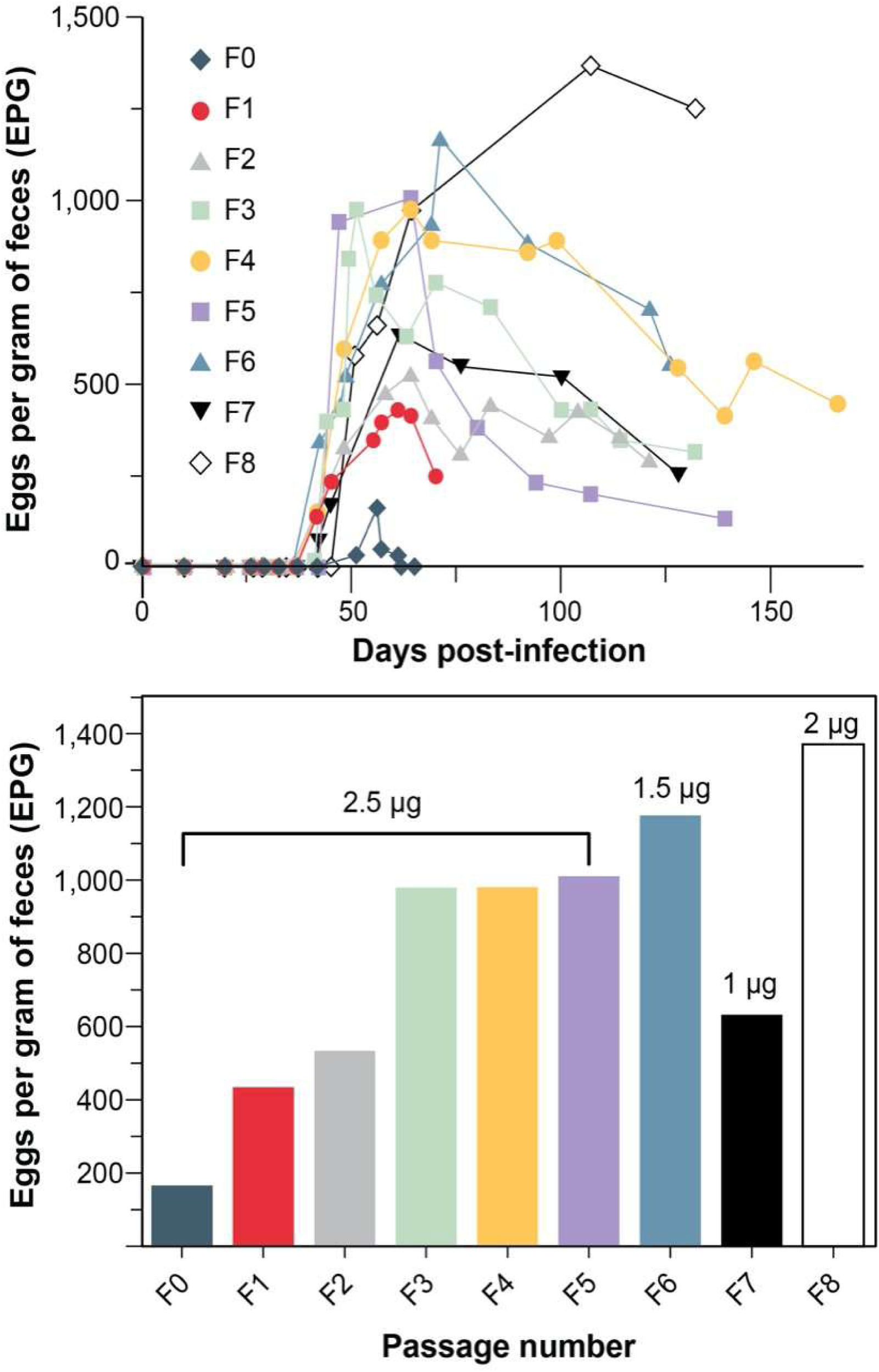
**Top**: Fecal egg excretion following infection of hamsters with Beposo strain of *N. americanus*. Data show serial eggs per gram (epg) of feces detected across the first 9 passages (F0-F8) in the animal model. **Bottom**: Highest fecal EPG recorded across the F0-F9 passages. Concentration (μg/ml) of dexamethasone in drinking water shown above bars.

Given the increasing egg output across the initial 5 passages, the concentration of dexamethasone in the drinking water was reduced to 1.5 µg/ml for the F6 passage. As shown in **Fig 1**, the peak egg excretion in the F6 infection (1172 EPG) was similar to the prior (F5) passage. However, when the dexamethasone was further reduced to 1.0 µg/ml for the F7 infection, there was a reduction in peak egg excretion to 627 EPG. Therefore, in order to sustain infection intensities needed to ensure propagation of the life cycle, the concentration of dexamethasone was increased to 2.0 µg/ml for the F8 infection, which exhibited a peak egg excretion of 1370 EPG. Overall, the longest duration of egg output recorded during this period was 166 DPI, which was noted in the F4 passage (**Fig 1**).

### Benzimidazole susceptibility of *Necator americanus* field isolates and targeted sequencing of partial beta-tubulin isotype 1

As shown in **Fig 2**, hookworm eggs from a laboratory strain of *A. ceylanicum* [18] and the newly adapted Beposo strain of *N. americanus* exhibited *in vitro* susceptibility to albendazole and mebendazole. Both species were more susceptible to albendazole than mebendazole, based on the egg hatch data. PCR amplification of genomic DNA from 19 adult worms from the initial passage did not identify 3 previously identified mutations in the beta-tubulin isotype 1 gene that have been associated with *in vitro* resistance to benzimidazole anthelminthics.

**Fig 2.**
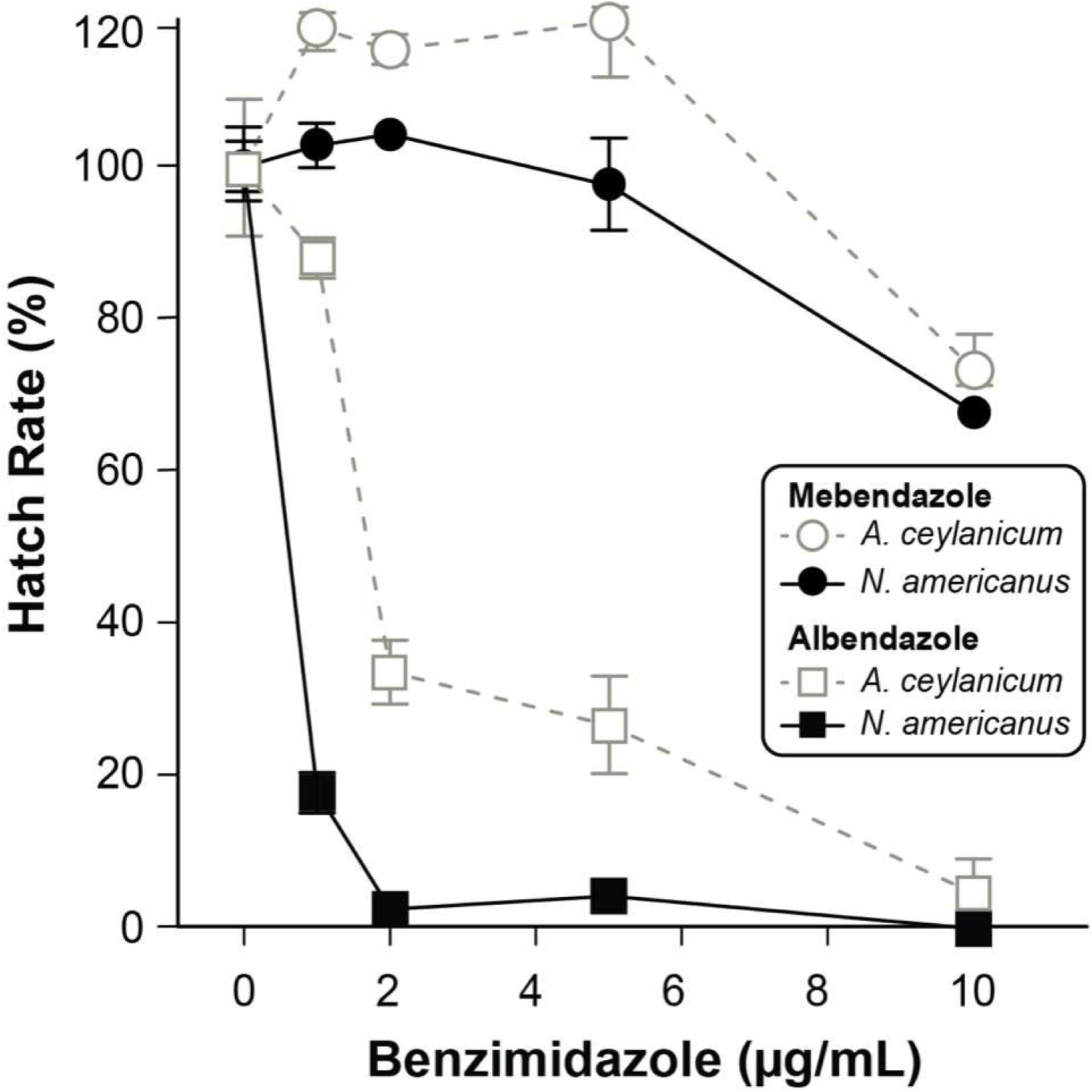
*In vitro* susceptibility of laboratory adapted hookworm strains to benzimidazole anthelminthics. Hookworm eggs isolated from hamsters infected with either *A. ceylanicum* (Ace) or the Beposo strain of *N. americanus* (Na) were incubated with increasing concentrations of mebendazole (Meb) or albendazole (Alb). Hatch rate was measured at 48 hours and compared to values recorded in the absence of drug.

### Phylogenetic analysis of partial *COX1* sequences from the Beposo strain

*COX1* sequences were successfully generated from 21 individual adult *N. americanus* F1 worms collected after the first passage of Beposo L3 in hamsters. Among these 21 individuals, 15 unique mitochondrial haplotypes were identified, differing from one another by 0.2–2.9% (1–16 bp along their 584 bp-aligned length). The final alignment for all partial *COX1* data was 622 base pairs in length, with 47 parsimony-informative sites, 59 singleton sites, and 516 invariant sites. Model selection as performed by ModelFinder in IQ-TREE indicated an HKY+F+I model as best-fit according to the Bayesian information criterion (BIC). Specimens from western Africa, including the specimens sequenced here from Ghana, the specimen from which the hybrid genome was assembled, and a single representative from Togo, formed a well-supported clade (UFBS=91) embedded among representative sequences from Brazil, China and Cambodia (**Fig 3**). GenBank accession numbers for all sequences included in the analysis, including those generated as part of this study, are provided in terminal labels in Fig 3. Among the individuals sequenced from Ghana, the specimen from which the hybrid genotype was assembled represented an additional 16^th^ unique *COX1* haplotype.

**Fig 3.**
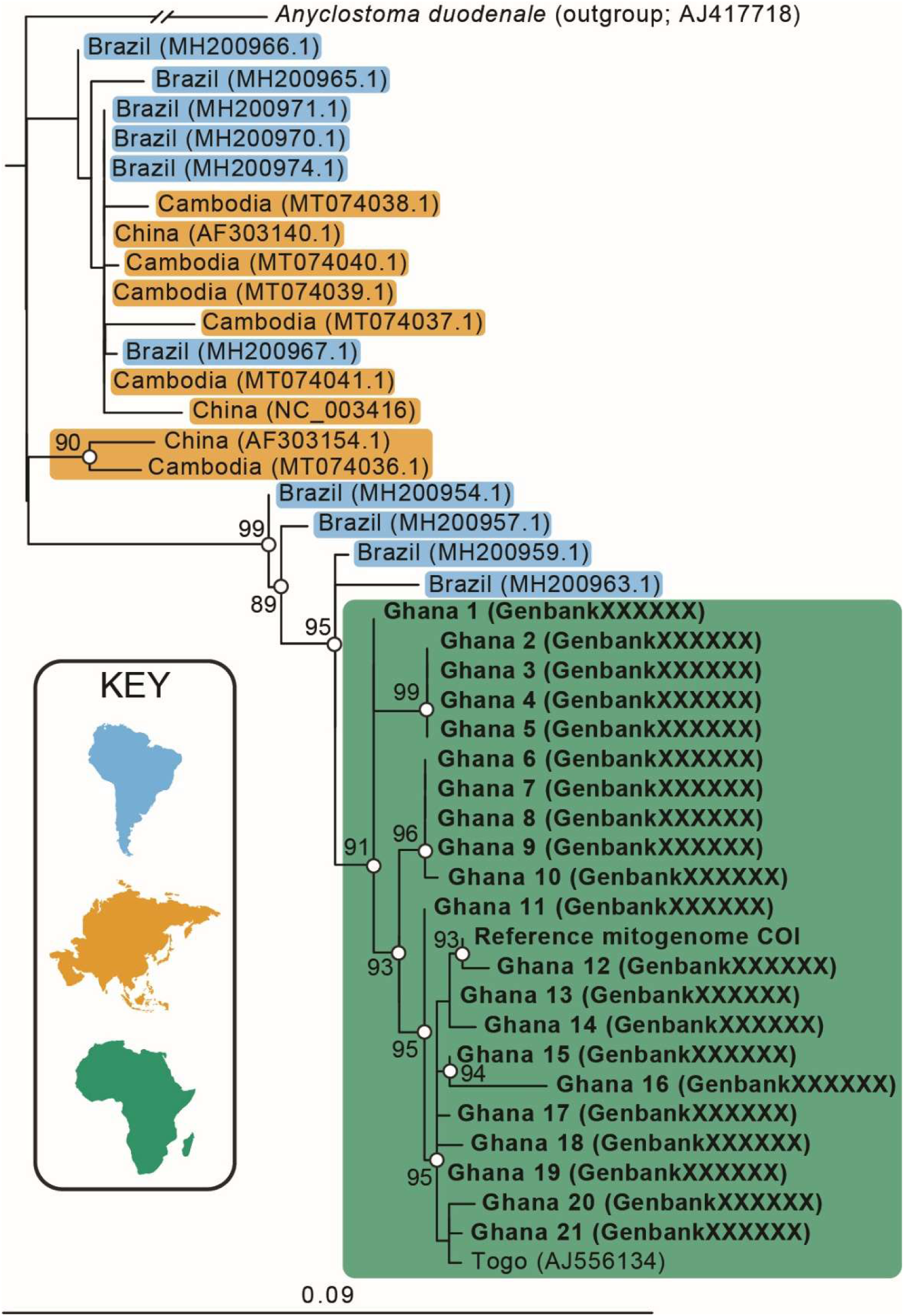
IQTree2 maximum likelihood topology for partial cytochrome c oxidase I subunit (COX1) from *Necator american*us. Bolded sequences are from individual Beposo strain adult worms (n=21) and the COX1 sequence extracted from the Beposo strain mitogenome assembly (n=1). Ingroup specimens, including those of *N. americanus* from other hookworm-endemic regions (n=20), are presented as collection country followed by GenBank accession number in parentheses. Topology is generated from a 622 base pair alignment specifying a HKY+F+I substitution model. Nodal support values are based on 1,000 ultrafast bootstrap replicates; support values <75 are not shown. A key to terminal coloration and a scale bar are presented at lower left; scale represents expected number of nucleotide substitutions per site. Submission to NCBI of accession numbers for newly reported COX1 sequences in process.

### Microsatellite analysis

Of the 40 microsatellite loci identified and evaluated on 20 F0 adult worms, 21 loci showed reliable amplification of a single band of the expected size. Diversity estimates for each locus are summarized in **Table 1** (summary statistics; full locus-level data in **S3 Table**) and show that all loci were polymorphic. The number of alleles per locus for the 20 F0 adult worms ranged from 3 to 16 (mean ± SD = 7.6 ± 3.7). Fifteen of the 21 loci showed statistically significant (p<0.05) deviations from Hardy-Weinberg equilibrium. Values of observed heterozygosity (Ho) ranged from 0.105 to 0.800 (mean ± SE = 0.465 ± 0.044) and expected heterozygosity (He) ranged from 0.370 to 0.901 (mean ± SE = 0.737 ± 0.029), respectively. Such deviations from Hardy-Weinberg equilibrium are commonly observed in parasite populations and may reflect population substructure, non-random mating, or sampling across multiple hosts. Observed inbreeding coefficient (Fis) ranged from −0.042 to 0.891 (mean ± SD = 0.327 ± 0.279).

**Table 1.** Characteristics of microsatellite loci of Beposo strain of *N. americanus*. Repeat motif, number of alleles (N), range of allelic size, observed (Ho) and expected (He) heterozygosity, Hardy–Weinberg P values (H–W) and inbreeding coefficient (Fis) are based on data from 20 adult worms.

| Locus | Repeat | Fragment Size (bp) | No of alleles | H<sub>o</sub><br>(observed) | H<sub>e</sub><br>(expected) | F<sub>is</sub> | H-W p-value |
| --- | --- | --- | --- | --- | --- | --- | --- |
| 257 | AC | 320-358 | 10 | 0.25 | 0.793 | 0.6345 | 0 |
| 1419 | AT | 222-242 | 9 | 0.5 | 0.727 | 0.1783 | 0.004 |
| 3087 | AATG | 204-306 | 10 | 0.556 | 0.86 | 0.2448 | 0 |
| 4783 | AAAT | 306-334 | 7 | 0.632 | 0.733 | 0.0645 | 0.202 |
| 5197 | AAC | 125-131 | 3 | 0.105 | 0.561 | 0.7986 | 0 |
| 5432 | ACGG | 106-138 | 5 | 0.579 | 0.596 | 0.1394 | 0.23 |
| 7291 | AC | 108-120 | 6 | 0.6 | 0.648 | 0.0338 | 0.176 |
| 10577 | AAC | 309-436 | 16 | 0.556 | 0.847 | 0.1153 | 0 |
| 10646 | AAC | 234-240 | 3 | 0.25 | 0.541 | 0.7795 | 0.004 |
| 15346 | AT | 230-254 | 10 | 0.4 | 0.858 | 0.6573 | 0 |
| 15719 | AG | 410-422 | 5 | 0.7 | 0.779 | 0.1444 | 0.244 |
| 15891 | AATG | 308-436 | 15 | 0.8 | 0.9 | 0.1118 | 0.014 |
| 16163 | AAAT | 100-226 | 12 | 0.611 | 0.901 | 0.3632 | 0 |
| 16559 | AAC | 219-231 | 5 | 0.55 | 0.559 | -0.0019 | 0.608 |
| 17420 | AAAT | 230-330 | 6 | 0.125 | 0.738 | 0.8913 | 0 |
| 17590 | ACC | 306-318 | 5 | 0.8 | 0.764 | -0.0421 | 0.979 |
| 18583 | ATCC | 98-128 | 8 | 0.611 | 0.826 | 0.2371 | 0.02 |
| 19084 | AAAT | 214-234 | 6 | 0.429 | 0.786 | 0.3307 | 0 |
| 19624 | AAAC | 222-334 | 9 | 0.316 | 0.798 | 0.412 | 0 |
| 21208 | AC | 204-218 | 3 | 0.154 | 0.37 | 0.4062 | 0.026 |
| 22283 | ACAT | 316-348 | 6 | 0.421 | 0.747 | 0.3611 | 0 |

The observed levels of expected heterozygosity (He 0.74) and allelic diversity are consistent with previous population genetic studies of *Necator americanus*, which have demonstrated substantial genetic variation within endemic populations and variability in diversity across geographic sites [55], supporting the interpretation that the Beposo strain represents a genetically diverse natural population.

Analyses carried out in MICRO-CHECKER 3.2.2 [56] indicated that 14 out of the 21 loci showed evidence for a potential null allele (**S4 Table**). There was no evidence for large allele dropout at any locus. Given the potentially high prevalence of null alleles at some loci, we used the adjusted genotypes as determined by MICRO-CHECKER, which corrects for the likely presence of null alleles, to test for HW equilibrium again at each locus. Correcting for null alleles resulted in 5 additional loci (1419, 5197, 10646, 19084, and 21208) no longer deviating significantly from Hardy-Weinberg equilibrium. However, most loci (N=10) still differed significantly (p<0.05) from Hardy-Weinberg equilibrium, even when using genotypes adjusted for null alleles.

### DNA extraction, library preparation, and next-generation sequencing

A total of 151ng of genomic DNA (gDNA) was extracted from the individual adult male *N. americanus* worm used for genome generation (**S5 Table**). From this extraction, 109ng of gDNA was used as input to a modified ONT MinION library preparation protocol designed to retain a higher percentage of DNA through the library preparation process. Based on results for other MinION libraries prepared from individual adult hookworms during protocol optimization, we anticipate that at 50-60% of this input was retained through our optimized library preparation approach (**S1 Fig**). The ONT MinION library was sequenced on two separate flow cells for a total of 37 hours resulting in 3.47 Gb of long-read data with a read length N50 of 6.3k base pairs (**S5 Table**, **S2 Fig**). A portion of the remaining gDNA (36 ng) was used for Illumina library preparation and sequencing, resulting in 12.61 Gb of short-read data. No significant contamination from host, bacteria, or human nucleic acid was identified in either read-level dataset (**S6 Table**).

### Genome and mitogenome assembly

K-mer analysis estimated the genome to be 202,800,477 bp with 2.21% heterozygosity (**Fig 4A-B**). The final hybrid assembly was 231,800,610 bp in length contained in 950 contigs, resulting in an assembly N50 of 449,410 (**Table 2**). The copy number spectrum plot confirmed that the assembly is heterozygous, and that duplication was successfully purged (**Fig 4C**). Genome wide GC content was 40.01%. miniBUSCO scores indicated that >95% of conserved nematode orthologs were identified in complete single copy (**Table 2**). Homology-based gene prediction through GeMoMa predicted 12,804 genes (**Table 2**). For MinION and Illumina read-level datasets, 84.54% and 87.98% of reads, respectively, mapped back to the final assembly, and no contamination was identified by BlobTools (**S3 Fig**). The mitogenome was 13,596 bp in length with a GC content of 23.3%, and is considered complete, with 12 coding sequences, large and small subunit rRNA, and 21 tRNAs identified (**Fig 5**).

**Fig 4.**
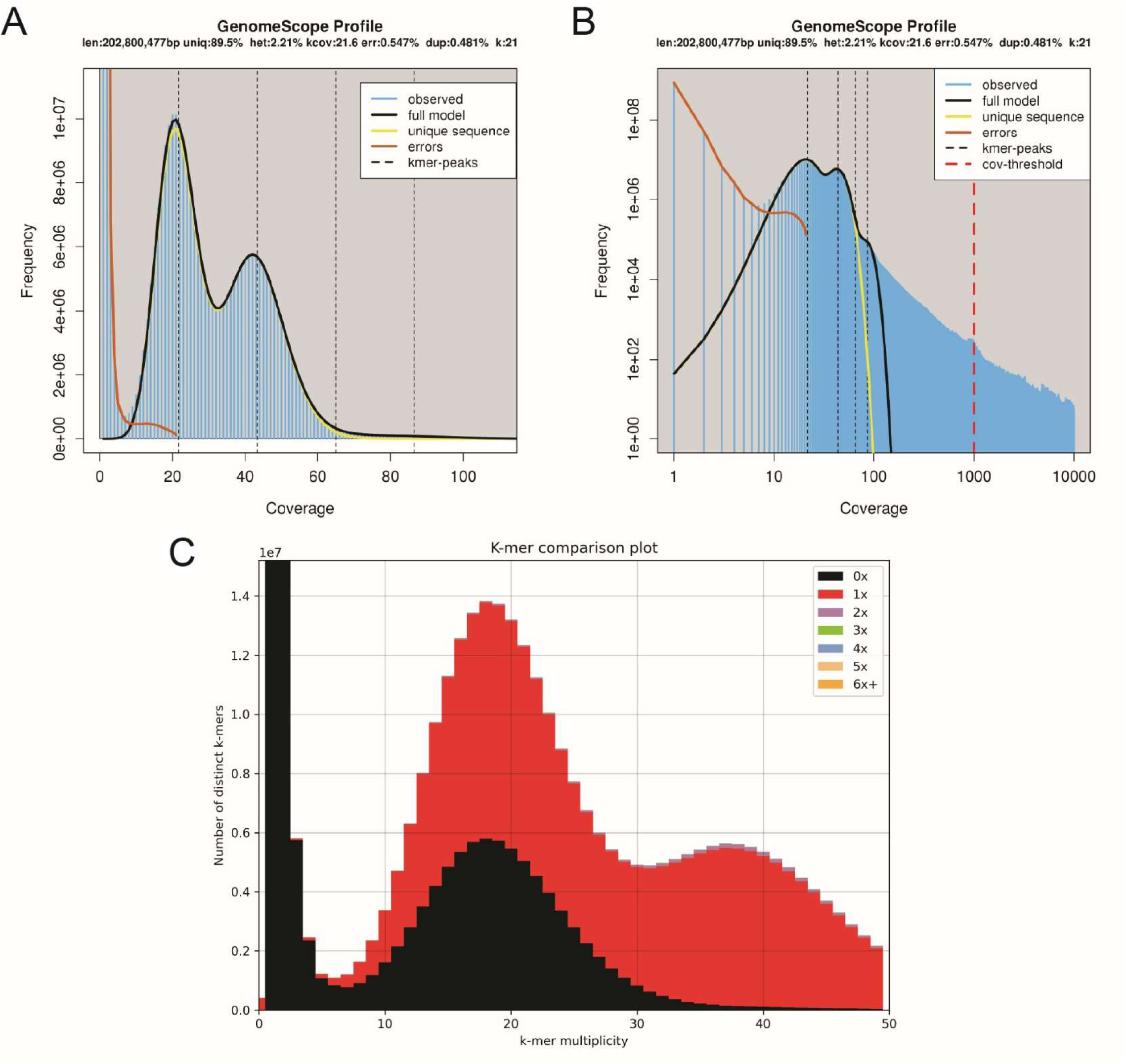
Focused (A) and complete (B) GenomeScope histograms generated from quality-controlled Illumina data and (C) K-mer multiplicity plot generated from purged hybrid assembly for *Necator americanus* from the Beposo strain.

**Fig 5.**
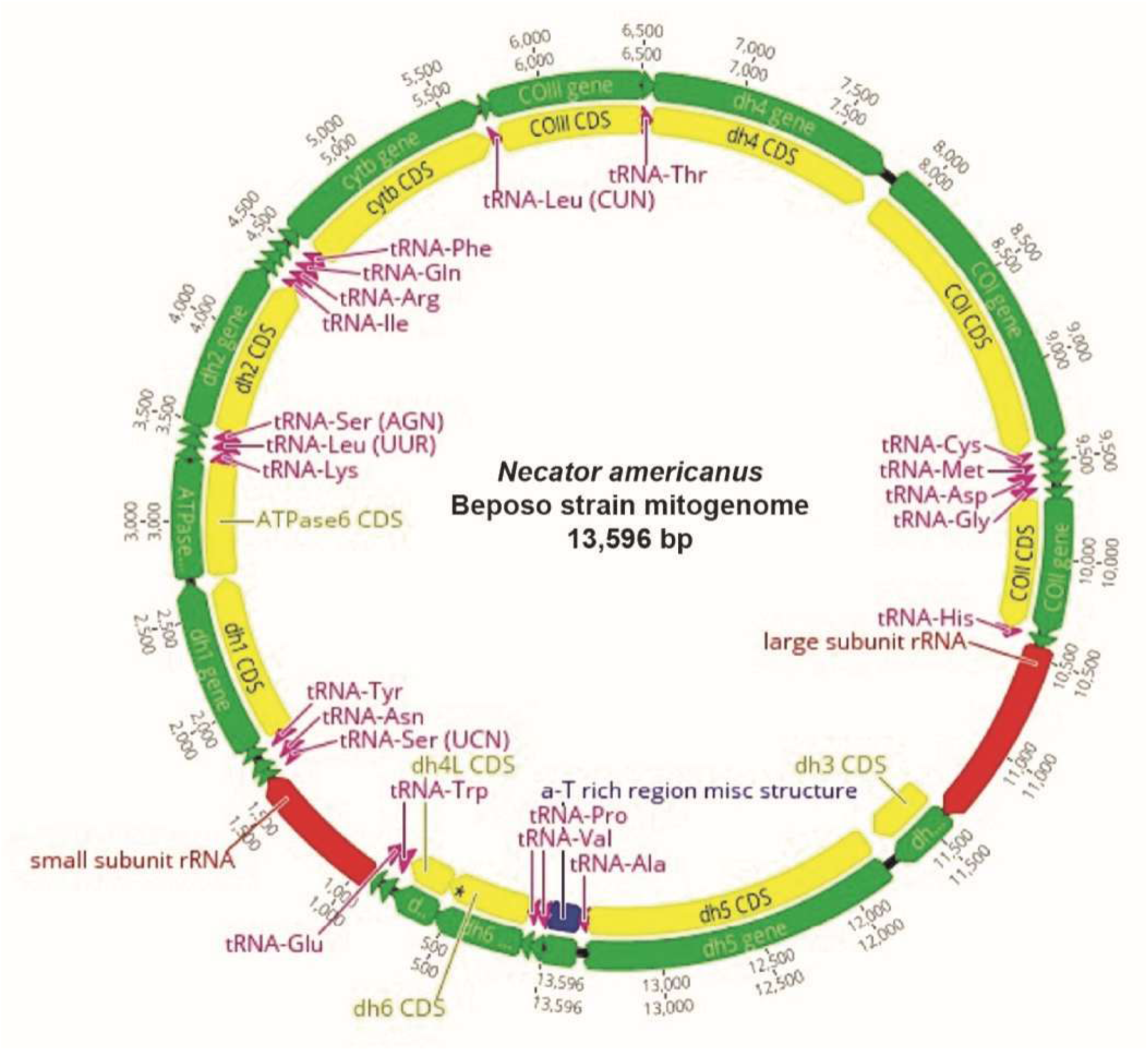
Graphical representation of the annotation of the mitochondrial genome from the Beposo strain assembly for *Necator americanus*.

**Table 2.**
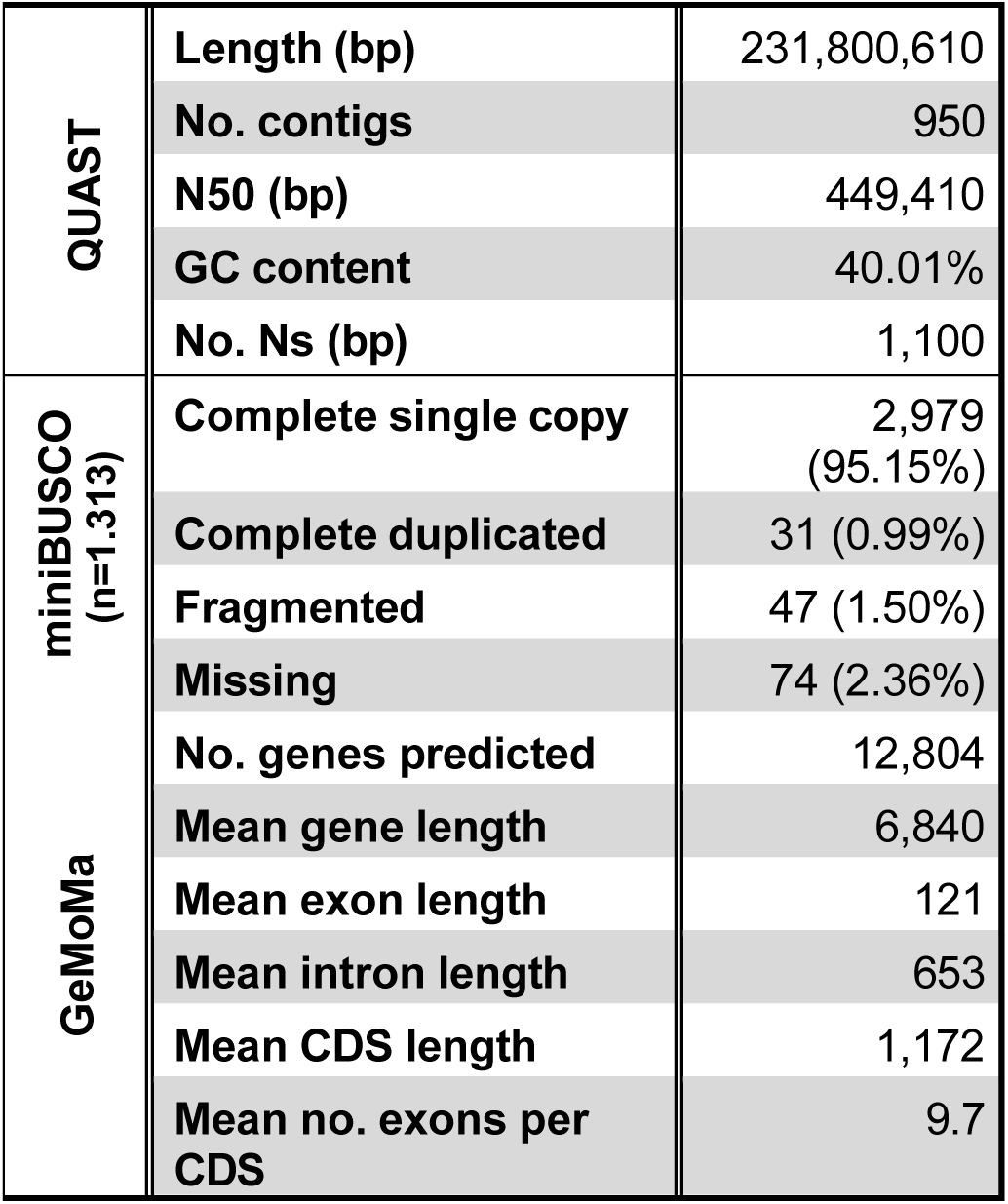
Quality metrics for the novel hybrid assembly of *Necator americanus*. For GeMoMa outputs, mean lengths are presented as number of base pairs. Abbreviations: bp=base pairs; CDS=coding sequence.

## Discussion

A major impediment to studying the biology and pathogenesis of human parasitic nematodes is the limited availability of reliable small animal models that can be maintained in a controlled laboratory setting. In the case of hookworm, the Golden Syrian hamster has been demonstrated to support the life cycle of human and animal hookworm species, including *N. americanus* and *A. ceylanicum* [13, 14, 17, 20, 21, 57, 58]. For both species, passage in the hamster has enabled the characterization of potential drugs and vaccines, as well as generation of stage specific parasite material for use in the development of diagnostics and identification of virulence factors.

As an obligate parasite, long term laboratory maintenance of the hookworm life cycle presents a particular challenge, requiring repeated passage and careful attention to the conditions necessary for consistent culturing of viable larvae from feces of infected animals. Initial passage of human field isolates is especially difficult, given that introduction of a human strain into a new host species is unpredictable and may not proceed at the level of infection intensity necessary for continued propagation, even in the most experienced hands. Perhaps for this reason, to date there have been few reports detailing the initial adaptation in the hamster model of *N. americanus* originally cultured from infected human study subjects [17, 20]. To our knowledge, this is the first description of an African strain of *N. americanus* that has been successfully passaged across multiple generations in a laboratory setting. Across the initial 9 passages (F0–F8) in the hamster, we observed a steady increase in peak egg excretion, demonstrating that maintenance of the Beposo strain is sustainable and exhibits a pattern consistent with stable adaptation from the human to rodent host.

As has been reported previously, successful completion of the *N. americanus* life cycle over initial passages has required the administration of glucocorticoids to recipient hamsters [17, 21]. In prior reports from China and India, it was demonstrated that exogenous steroid treatment of the hamsters could eventually be reduced or withdrawn. At least through the first 9 passages of the Beposo strain, we have not been able to reliably obtain viable larvae necessary for continued passage without providing the hamsters dexamethasone in drinking water. However, reducing the concentration of steroid from 2.5 µg/ml (F0) to a dose of 2.0 µg/ml (F8) showed comparable levels of fecal egg excretion and adult worm recovery. Subsequent passages (F9-F20) have enabled further reductions in glucocorticoid dosing of recipient hamsters, which is currently maintained at 1.0 µg/ml for routine maintenance of the *N. americanus* life cycle in the laboratory.

In order to characterize the newly adapted *N. americanus* strain, we utilized an egg hatch assay that allows for *in vitro* screening of isolates for susceptibility to anthelminthics [35, 59, 60]. Hatching of eggs purified from hamster feces was inhibited in the presence of increasing concentrations of albendazole (**Fig 2**). The Beposo strain was found to be more sensitive to albendazole than mebendazole, which is consistent with the differential susceptibility to these related benzimidazoles observed previously for *N. americanus* [61–63]. To further assess the potential for drug resistance-associated mutations in the Beposo strain, we amplified regions of genomic DNA corresponding to the 3 well described single nucleotide polymorphisms (SNPs) in the beta-tubulin isotype 1 gene that are thought to confer benzimidazole resistance. Based on Sanger sequencing of PCR amplicons, we did not find evidence of known resistant genotypes in 20 individual adult *N. americanus* worms from the initial (F0) passage. This result, although representing a small sample of the hookworm population in Beposo, is consistent with prior deep sequencing data from hookworm eggs previously collected in Kpandai District in Ghana [64]. At present, it is unknown what additional selective pressure on the hookworm genome may be introduced by sequential passage through the hamster host.

Analysis of partial *COX1* sequence data indicates genetic polymorphisms that distinguish *N. americanus* in Ghana from other hookworm endemic regions, including Brazil, China and Cambodia, but that are more genetically similar to sequences from the adjacent country of Togo (**Fig 3**). The specimens sequenced here are among the first *COX1* sequences generated from *N. americanus* from Ghana, significantly expanding the number of sequences publicly available for *N. americanus* from Africa.

The microsatellite analysis provides important population-level context for the Beposo strain. The observed levels of allelic diversity (mean 7.6 alleles per locus) and heterozygosity (He 0.74) are consistent with prior microsatellite-based analyses of hookworms and other parasitic nematodes, which have demonstrated substantial genetic variation within endemic populations [55]. Such patterns are generally attributed to large effective population sizes and ongoing transmission in natural settings. Deviations from Hardy-Weinberg equilibrium at multiple loci, even after correction for null alleles, are also commonly reported in microsatellite datasets for parasitic helminths and may reflect population substructure or sampling from multiple hosts. The presence of null alleles at several loci is not unexpected and does not alter the overall conclusion that the founding Beposo parasite population was genetically heterogeneous.

Using additional ONT MinION libraries generated for single hookworms, we present here for the first time comparative data to demonstrate that this approach represents a substantial improvement in gDNA retention through the ONT MinION library preparation process (**S1 Fig**). The optimized library preparation approach, which largely consists of extended bead binding and elution times and titrating ONT Long and Short Fragment Wash Buffers allows for retention of nearly 60% of input gDNA, as compared to the standard protocol in which less than 4% of input is retained (**S1 Fig**).

We also detail a contiguous and complete whole genome assembly for *N. americanus* generated from a single first passage adult male from the newly established Beposo strain. The final assembly is 202.8Mb in length, is relatively contiguous with an N50 >449k, and is considered highly complete with >95% complete single copy nematode orthologs identified. To our knowledge, this assembly represents the first for *N. americanus* from an African strain, and the first generated from a single adult worm. It joins two other published whole genome assemblies for the species: that of Tang et al [53] and improved by Logan et al [65], generated from worms from the Anhui strain isolated from Hunan Province, China (GenBank no. GCA_000507365.1), and the chromosome-scale annotated assembly generated from a pool of adult male and female worms that were presumably isolated from Brazil (GenBank no. GCA_031761385.1). While the Beposo strain assembly represents an important genomic resource, it should be considered a draft assembly at this stage. Additional data types are needed to complete it, including chromosome contact information (such as Hi-C sequencing) to scaffold the assembly into chromosomes, and RNA sequencing data from multiple life-cycle stages to improve the gene annotation. Efforts to generate and incorporate these data types are in progress.

While the genome assembly presented here provides a high-quality reference derived from a single adult worm, it represents only a single realization of the genetic makeup of the parasite population. As such, it does not capture the extent of standing genetic variation present in the original field isolate. Complementary microsatellite analysis addresses this limitation by providing a population-level view of genetic diversity among parasites collected from Beposo, Ghana. Across multiple loci, the presence of several alleles per locus and moderate-to-high levels of heterozygosity demonstrate that the founding population was genetically heterogeneous rather than clonal. This distinction is important for interpreting the laboratory adaptation process, as it suggests that serial passage in hamsters was initiated from a diverse founding population, allowing for potential selection and adaptation across generations rather than expansion of a single genotype. This may help explain the progressive increase in infection intensity observed across passages, consistent with selection acting on standing genetic variation within the founding population.

In this context, the genome and microsatellite data (**Table 1 and S3 Table**) provide complementary and non-redundant insights: the genome assembly establishes a high-quality reference framework, while microsatellite markers describe population structure and genetic diversity in the first generation of the newly established model. Importantly, these population-level insights cannot be inferred from a single-worm genome assembly, underscoring the non-redundant value of the microsatellite data. Together, these data provide a more complete representation of the genetic landscape of the Beposo strain and establish a baseline for future studies aimed at tracking genetic changes during laboratory passage, drug exposure, or other selective pressures. This is particularly important for hookworm, where population-level genetic diversity may influence transmission dynamics, drug response, and the emergence and spread of resistance in endemic settings. Although some microsatellite loci showed evidence of null alleles and deviations from Hardy-Weinberg equilibrium, these patterns are not unexpected in small, structured populations and do not detract from the overall conclusion of substantial genetic diversity.

Regionally specific reference genomes for helminths, like the Beposo strain assembly presented here, are of great value in experimental, evolutionary, and comparative studies [66, 67]. This is especially true for those pathogens, including hookworm and other STHs, for which the primary method of control is targeted mass drug administration [68]. Infection with *N. americanus* remains prevalent in sub-Saharan Africa, and at the recommendation of the WHO, many countries have sustained national deworming efforts to reduce morbidity and curb transmission [1, 69]. However, because widespread anthelminthic use has the potential to impose selective pressure in favor of drug tolerance, there is a need for resources and approaches to detect and characterize genetically mediated resistance at its first emergence [64, 70–76]. Ultimately, access to parasite proteins and individual life cycle stages from the newly adapted strain of *N. americanus* will create opportunities to identify virulence factors and gene expression patterns, which will further advance understanding of hookworm biology. In addition, a reproducible small animal model that utilizes the most common human hookworm worldwide provides a platform for pathogenesis studies and preclinical development of new drugs, diagnostics and vaccines. Finally, the Beposo strain genome assembly has the capacity to aid in identifying genomic patterns associated with hookworm evolution and adaption to the hamster host, as well as tracking changes in parasite populations and monitoring mutations associated with reduced treatment efficacy in endemic communities repeatedly exposed to anthelminthics.

## Acknowledgements

The authors would like to thank the people of Beposo, Ghana for their participation in this research study. We would also like to acknowledge Mr. Frimpong Mensah for logistics and transportation support. This work was completed utilizing the Holland Computing Center of the University of Nebraska, which receives support from the UNL Office of Research and Innovation, and the Nebraska Research Initiative. Additional support was provided by Yale University through a Lindsay Fellowship for Research in Africa (MacMillan Center for International and Area Studies) and the Yale Institute for Biospheric Studies.

---

**S1 Fig.**
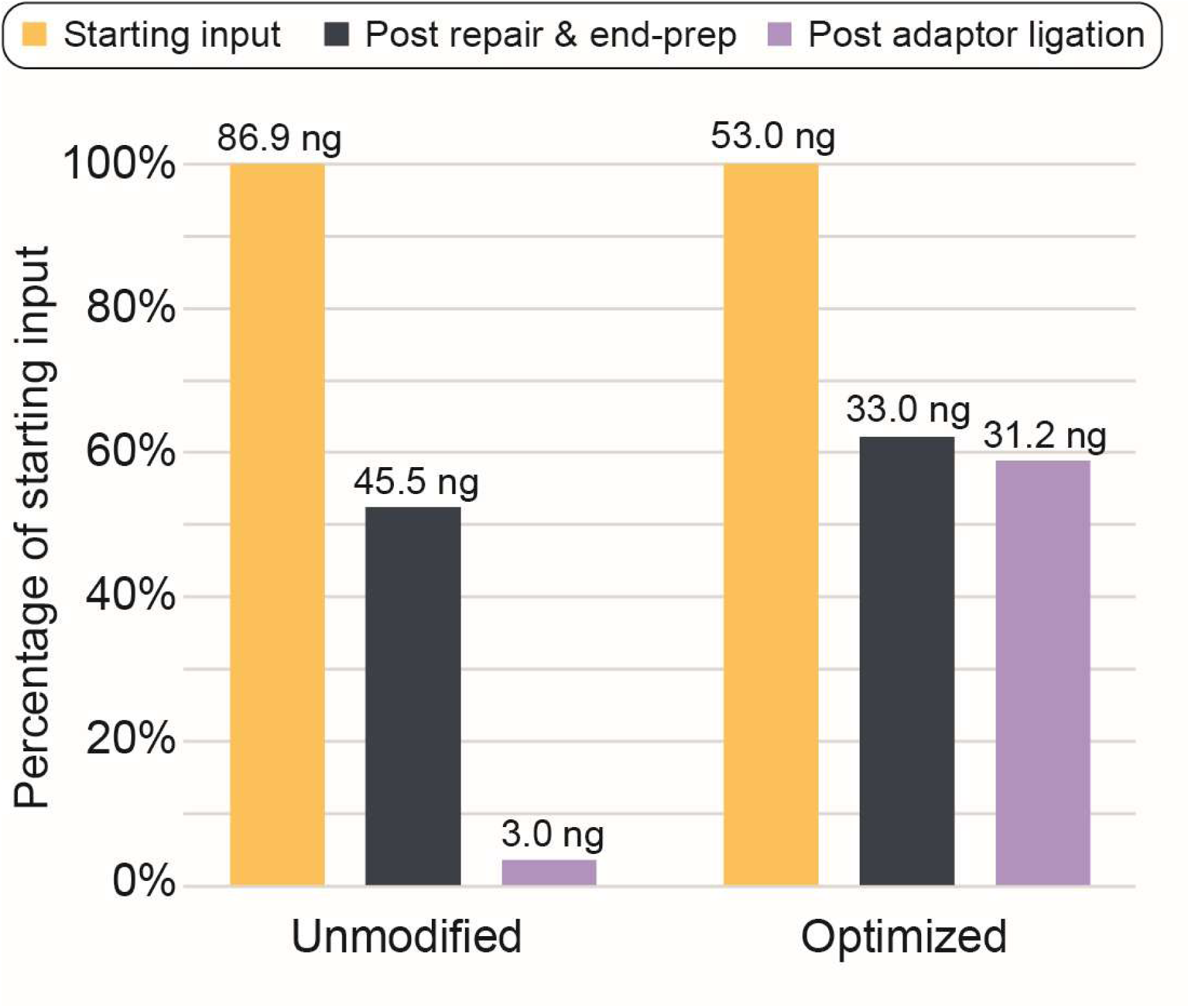
Results of optimized Oxford Nanopore Technologies MinION library preparation protocol demonstrating greater retention of genomic DNA from individual adult males of Beposo strain *Necator americanus*.

**S2 Fig.**
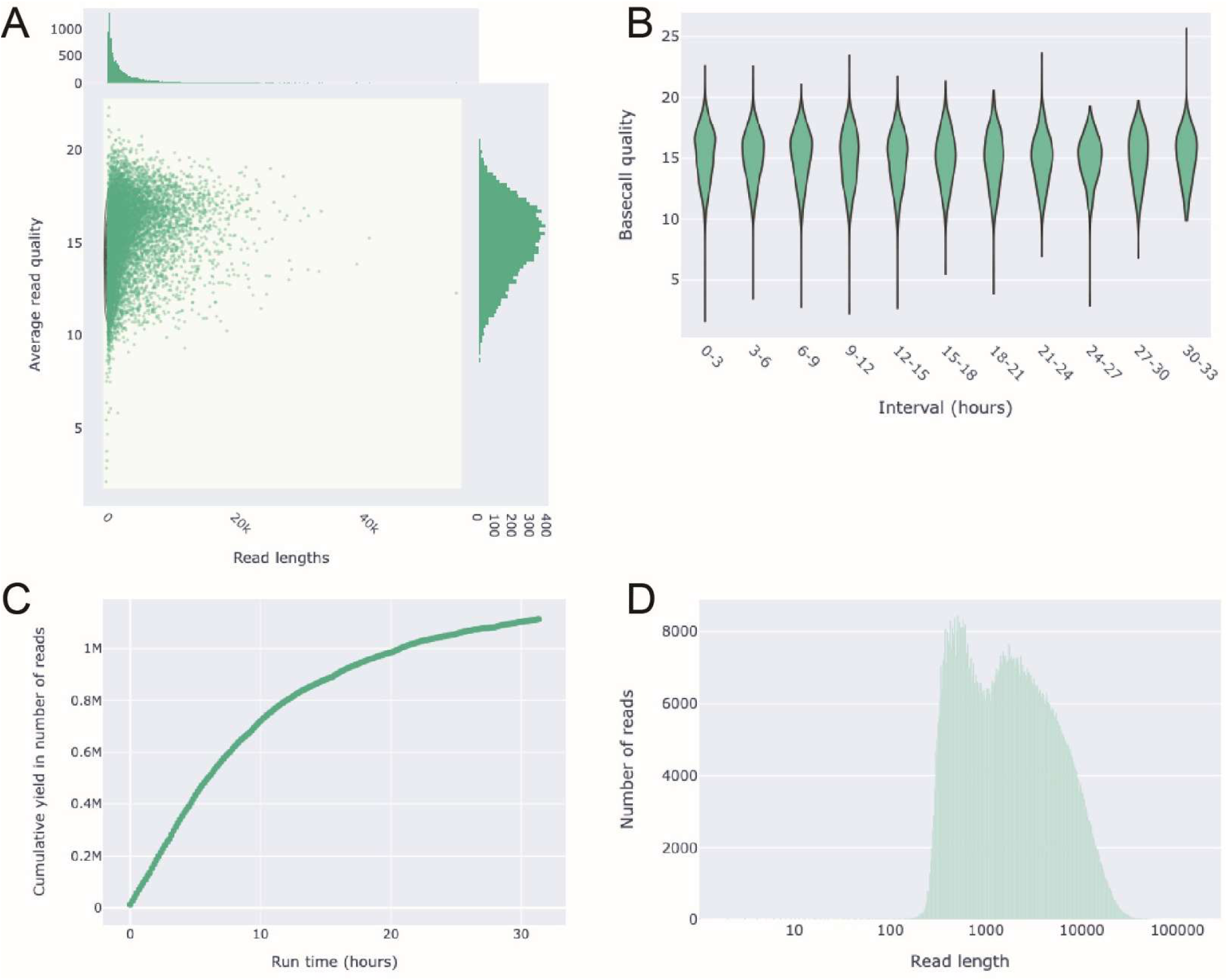
Output from NanoPlot summarizing the combined quality and efficiency of the two MinION sequencing runs for Beposo strain *Necator americanus*. (A) Histogram of read length versus average read quality score. (B) Range of read quality scores versus time. (C) Cumulative yield plot. (D) Non-weighted log-transformed histogram of read lengths.

**S3 Fig.**
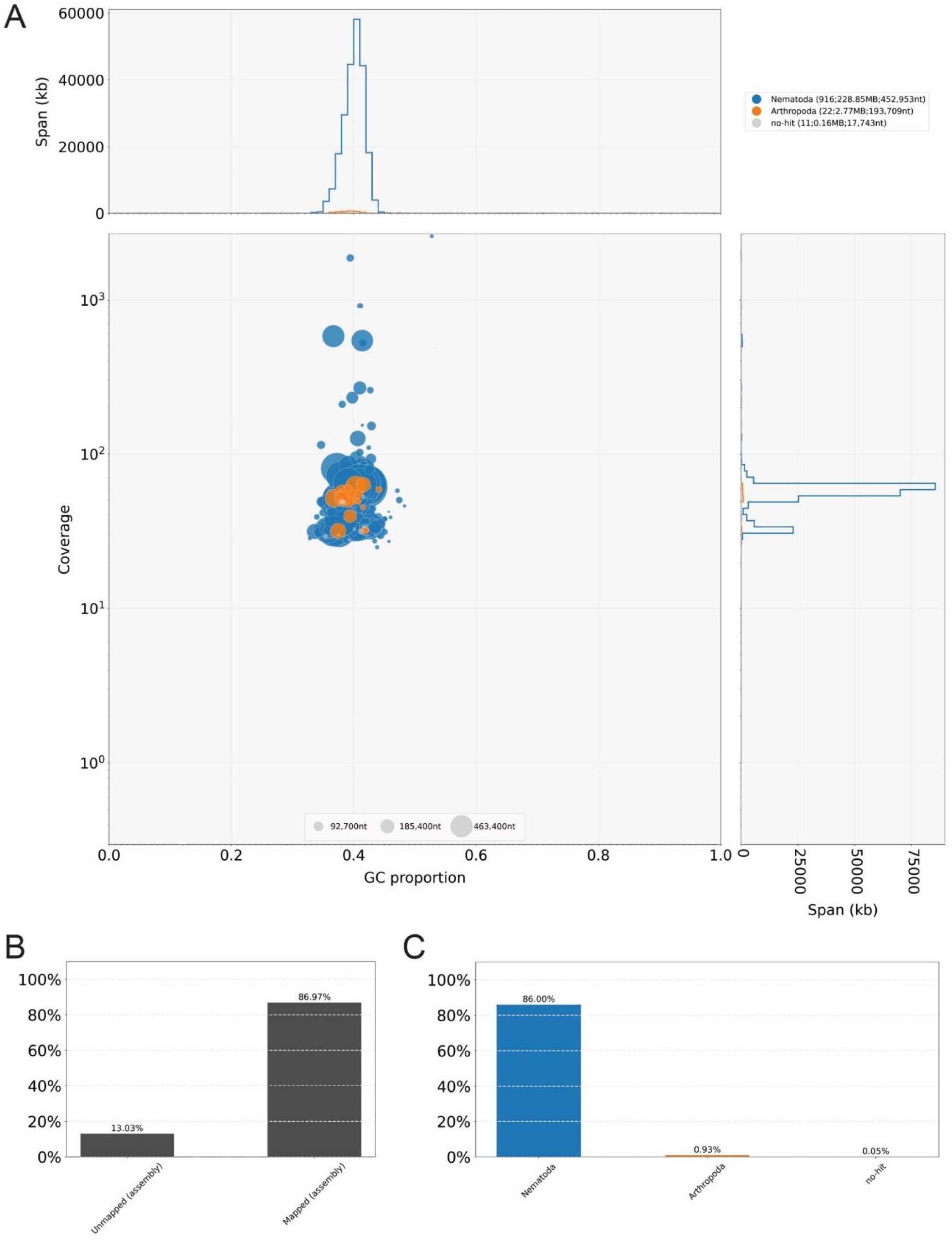
BlobTools output the Beposo strain assembly for *Necator americanus*. (A) BlobPlot. (B–C) Read coverage plots.

**S1 Table.**
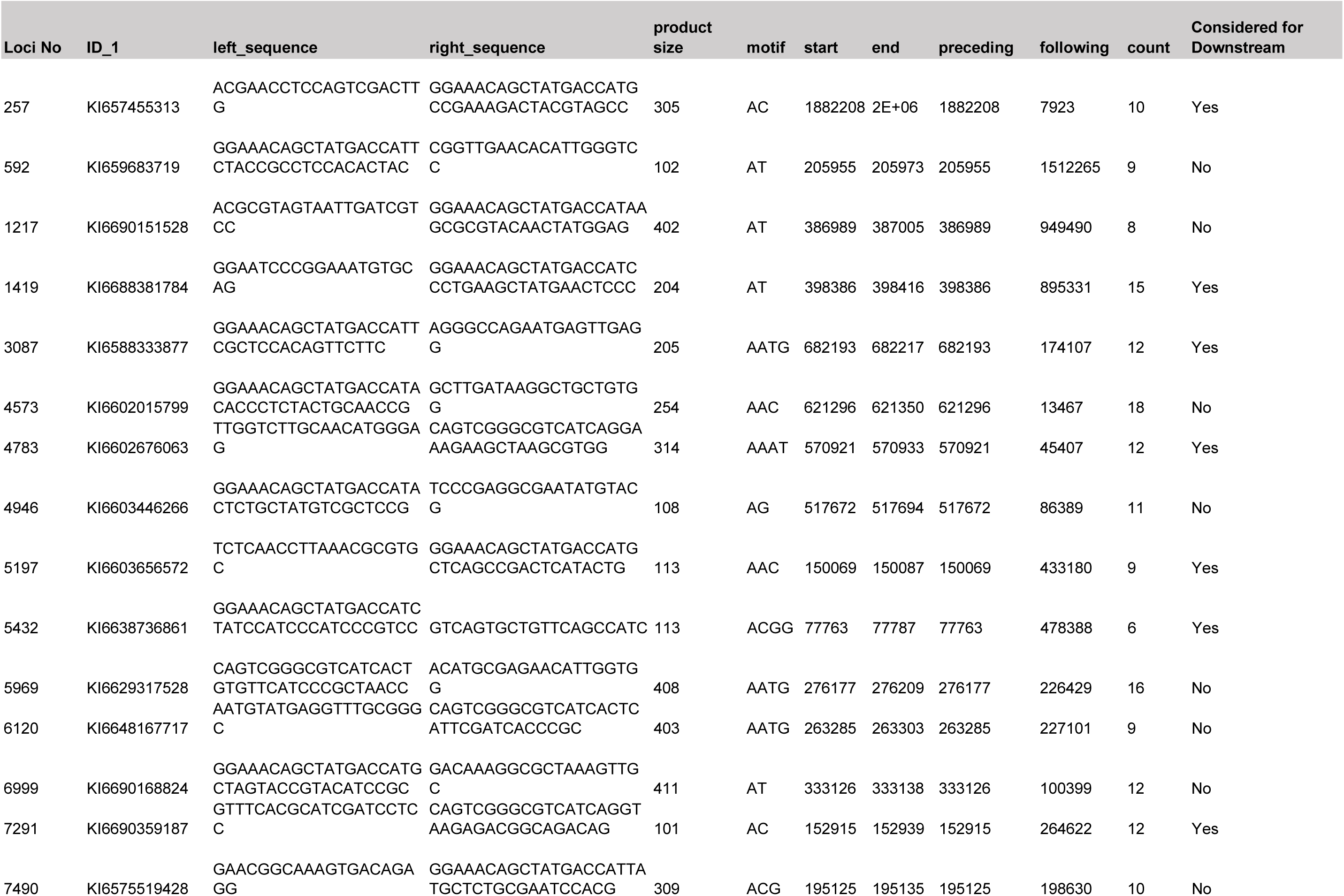
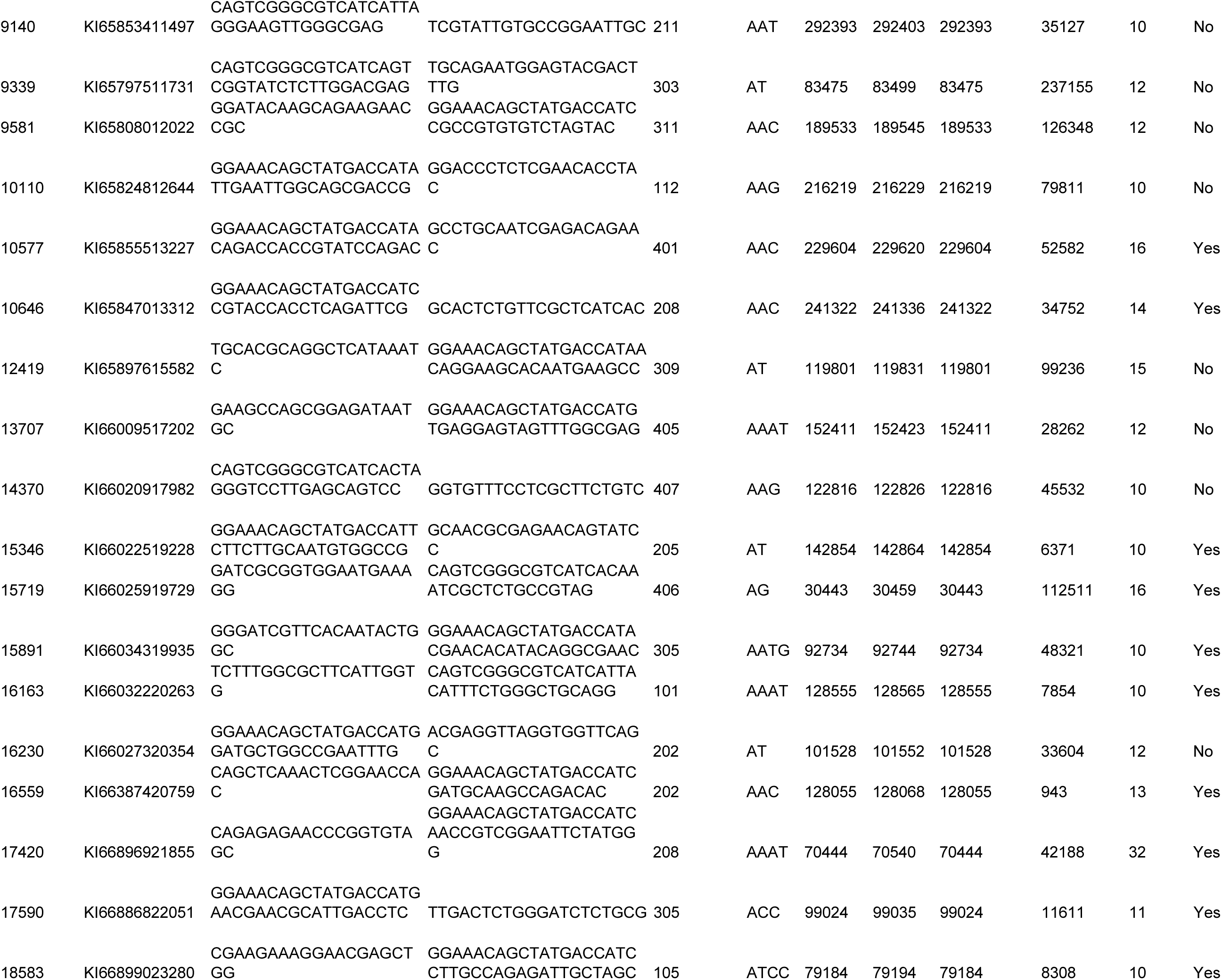
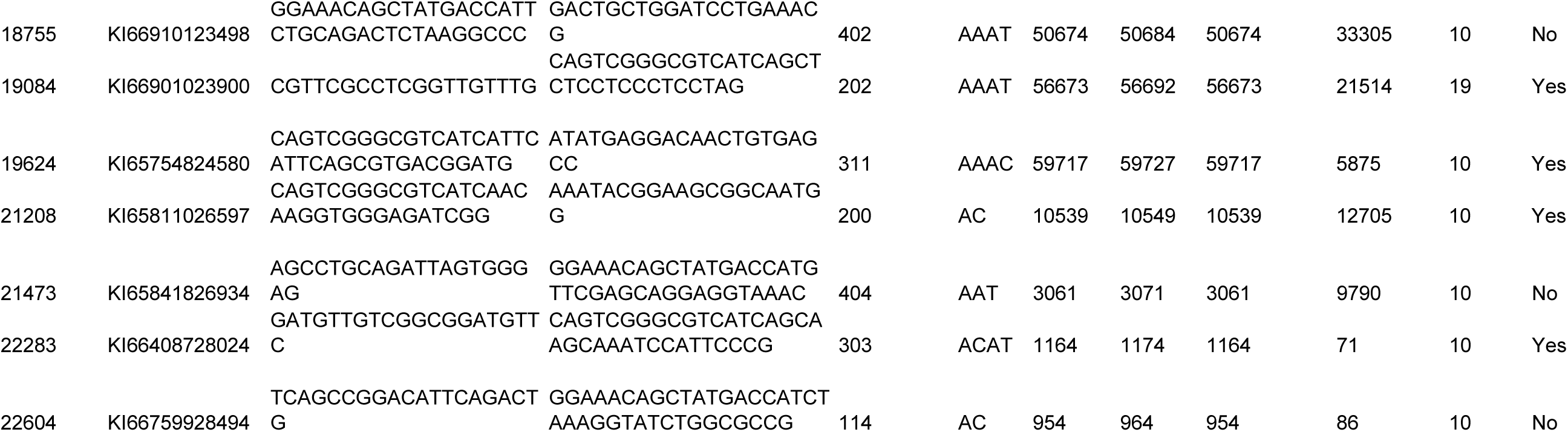
The 40 microsatellite loci from Necator americanus (PRJNA72135) that were shorlisted from the complete genome obtained from wormbase.org. A total of 22,632 microsatellite regions were identified and a subset of 40 microsatellites were shortlisted based on varying repeat sequences and fragment sizes

**S2 Table.**
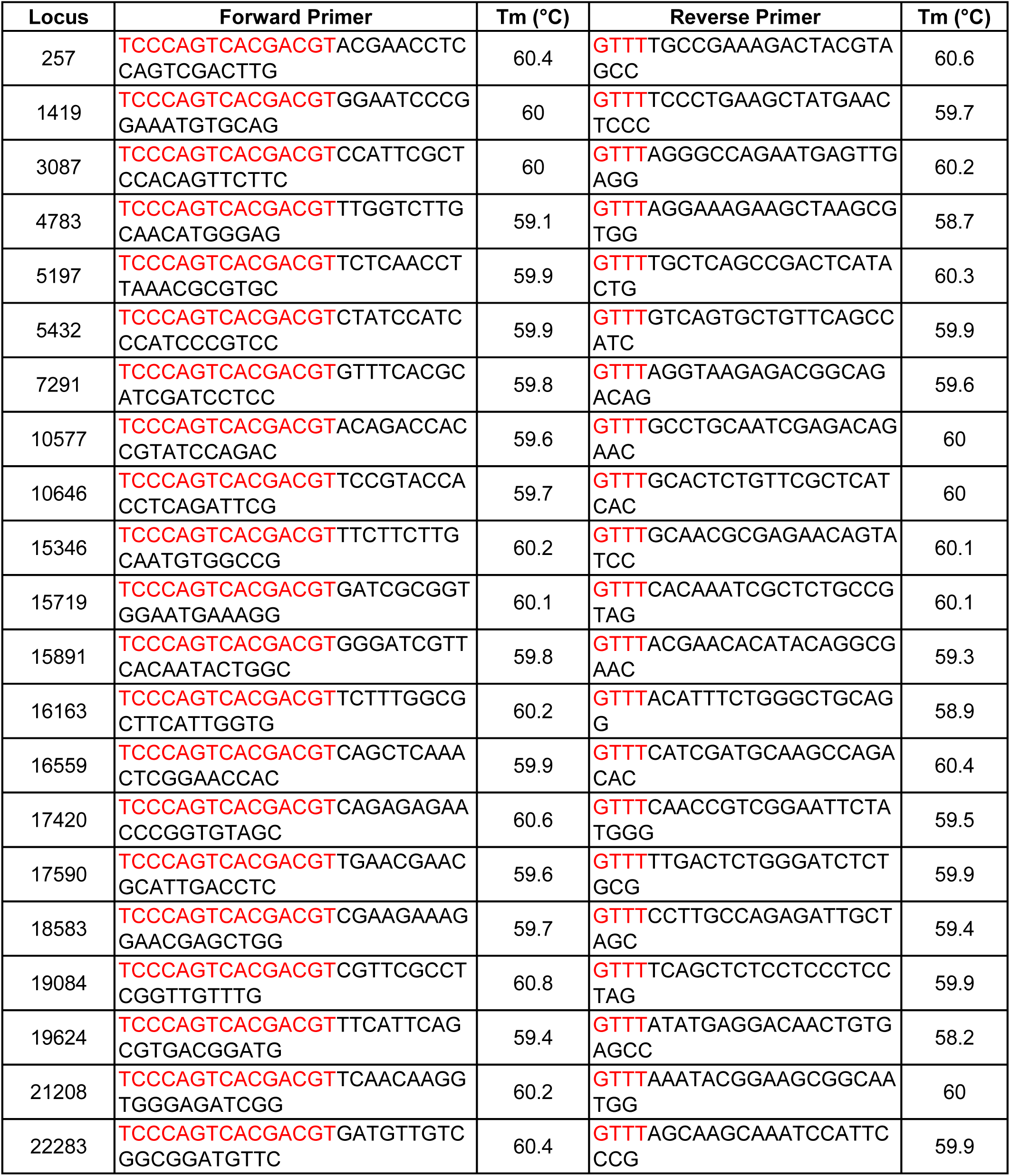
Primer sequences and annealing temperatures of 21 di, tri and tetra microsatellite loci for N. americanus (BioProject PRJNA72135). Forward primers were tagged with M13 universal sequence (in red). Reverse primers are tagged with GTTT pigtail (also in red).

**S3 Table.**
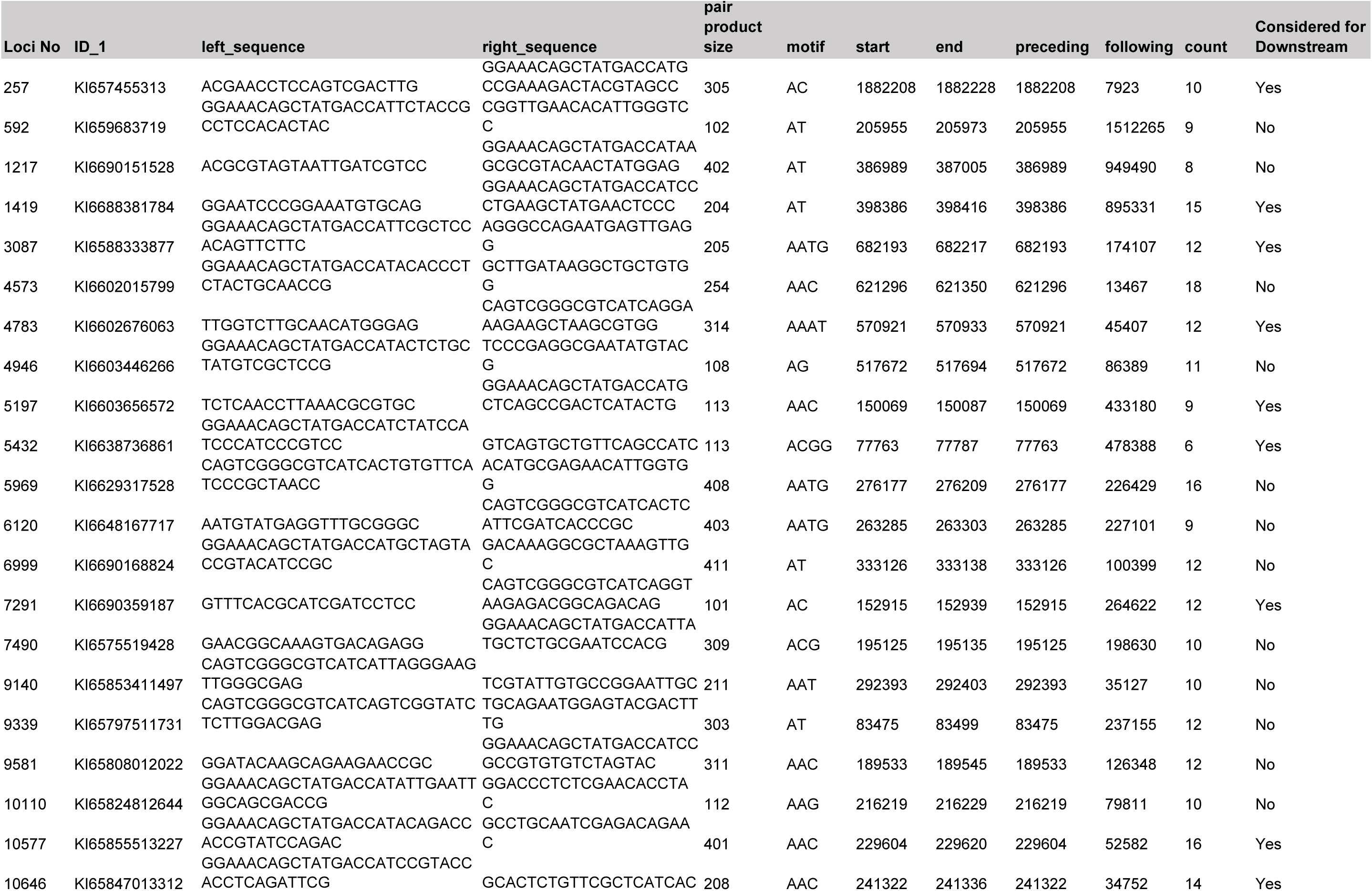
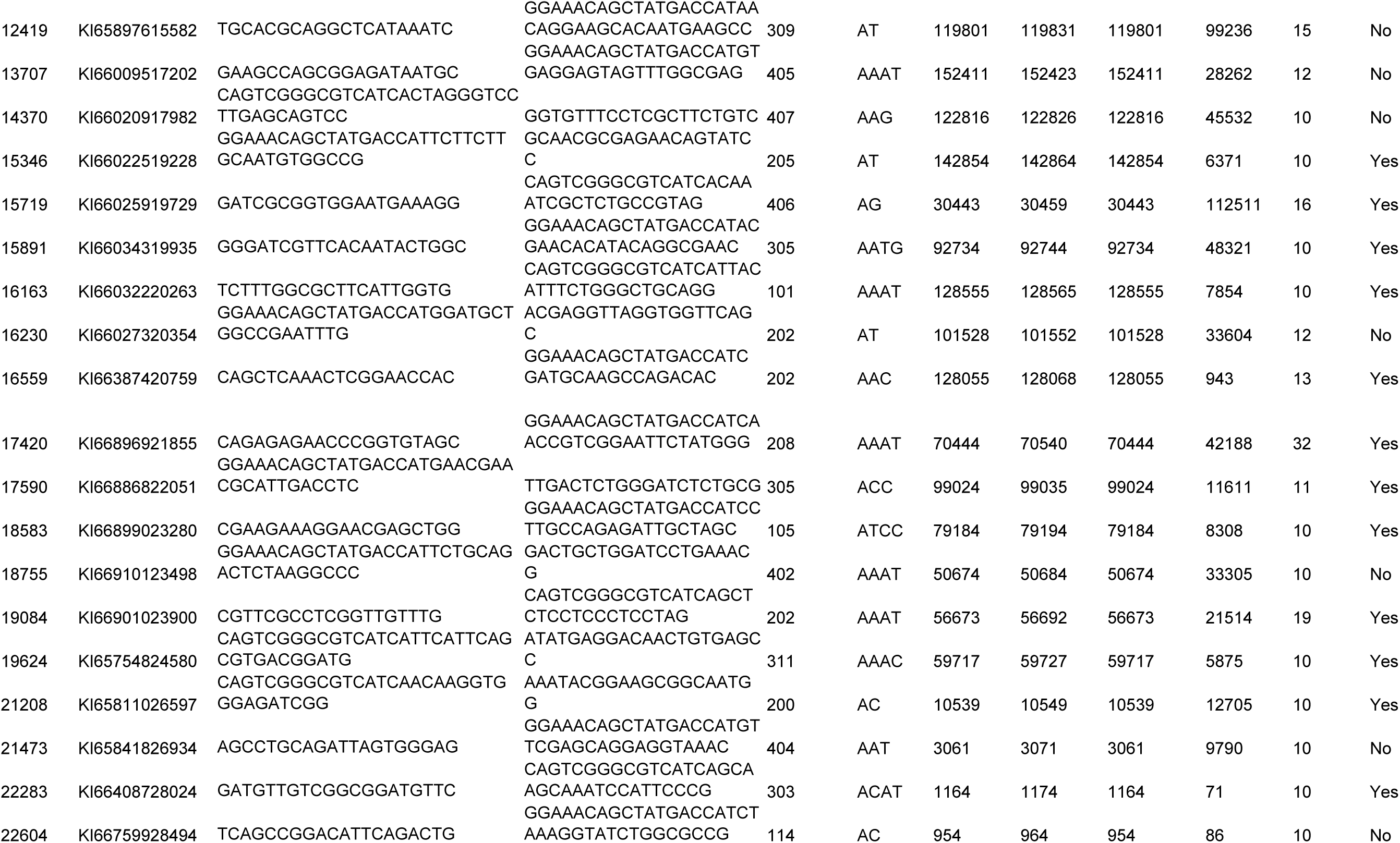
The 40 microsatellite loci from *Necator americanus* (PRJNA72135) that were shorlisted from the complete genome obtained from wormbase.org. A total of 22,632 microsatellite regions were identified and a subset of 40 microsatellites were shortlisted based on varying repeat sequences and fragment sizes.

**S4 Table.** Results from MICRO-CHECKER version 2.2.3. Several loci show evidence for a null allele. There was no evidence for ‘Large allele dropout’ at any locus. Results are consistent with this population possibly being in Hardy Weinberg equilibrium with loci 257, 1419, 3087, 5197, 9140, 10577, 10646, 15346, 16163, 17420, 18583, 19084, 19624, 21208, 22283, showing signs of a null allele. MicroChecker flagged 3 loci as having potential evidence of errors due to stuttering: 5197, 10646, 22283. These loci were scored independently by two different observers, then re-scored again blindly by one observer who paid specific attention to stutter peaks that may have been masking alleles. No changes to the allele calls were deemed necessary.

| **Locus** | **Null Present** | **Oosterhout** | **Chakraborty** | **Brookfield 1** | **Brookfield** |
| --- | --- | --- | --- | --- | --- |
| **257** | **yes** | **0.321** | **0.5204** | **0.3026** | **0.3026** |
| **1419** | **yes** | **0.1724** | **0.1849** | **0.1314** | **0.301** |
| **3087** | **yes** | **0.1786** | **0.2148** | **0.1635** | **0.3143** |
| 4783 | no | 0.0712 | 0.0741 | 0.0584 | 0.1699 |
| **5197** | **yes** | **0.3557** | **0.684** | **0.2919** | **0.3722** |
| 5432 | no | -0.0049 | 0.0142 | 0.0104 | 0.153 |
| 7291 | no | 0.0497 | 0.0381 | 0.0288 | 0.0288 |
| **9140** | **yes** | **0.3252** | **0.497** | **0.31** | **0.4875** |
| **10577** | **yes** | **0.1822** | **0.2079** | **0.1579** | **0.3107** |
| **10646** | **yes** | **0.2371** | **0.3681** | **0.189** | **0.189** |
| **15346** | **yes** | **0.2531** | **0.3638** | **0.2463** | **0.2463** |
| 15719 | no | 0.0425 | 0.0533 | 0.0443 | 0.0443 |
| 15891 | no | 0.0535 | 0.0588 | 0.0526 | 0.0526 |
| **16163** | **yes** | **0.1561** | **0.1918** | **0.1526** | **0.3013** |
| 16559 | no | 0.0128 | 0.0079 | 0.0056 | 0.0056 |
| **17420** | **yes** | **0.3885** | **0.7104** | **0.3528** | **0.5714** |
| 17590 | no | -0.0217 | -0.0232 | -0.0206 | 0 |
| **18583** | **yes** | **0.1293** | **0.1493** | **0.1175** | **0.2795** |
| **19084** | **yes** | **0.2246** | **0.2941** | **0.2** | **0.5616** |
| **19624** | **yes** | **0.2974** | **0.4328** | **0.2681** | **0.3424** |
| **21208** | **yes** | **0.2301** | **0.4124** | **0.1577** | **0.6147** |
| **22283** | **yes** | **0.2164** | **0.2788** | **0.1864** | **0.2737** |

**S5 Table.**
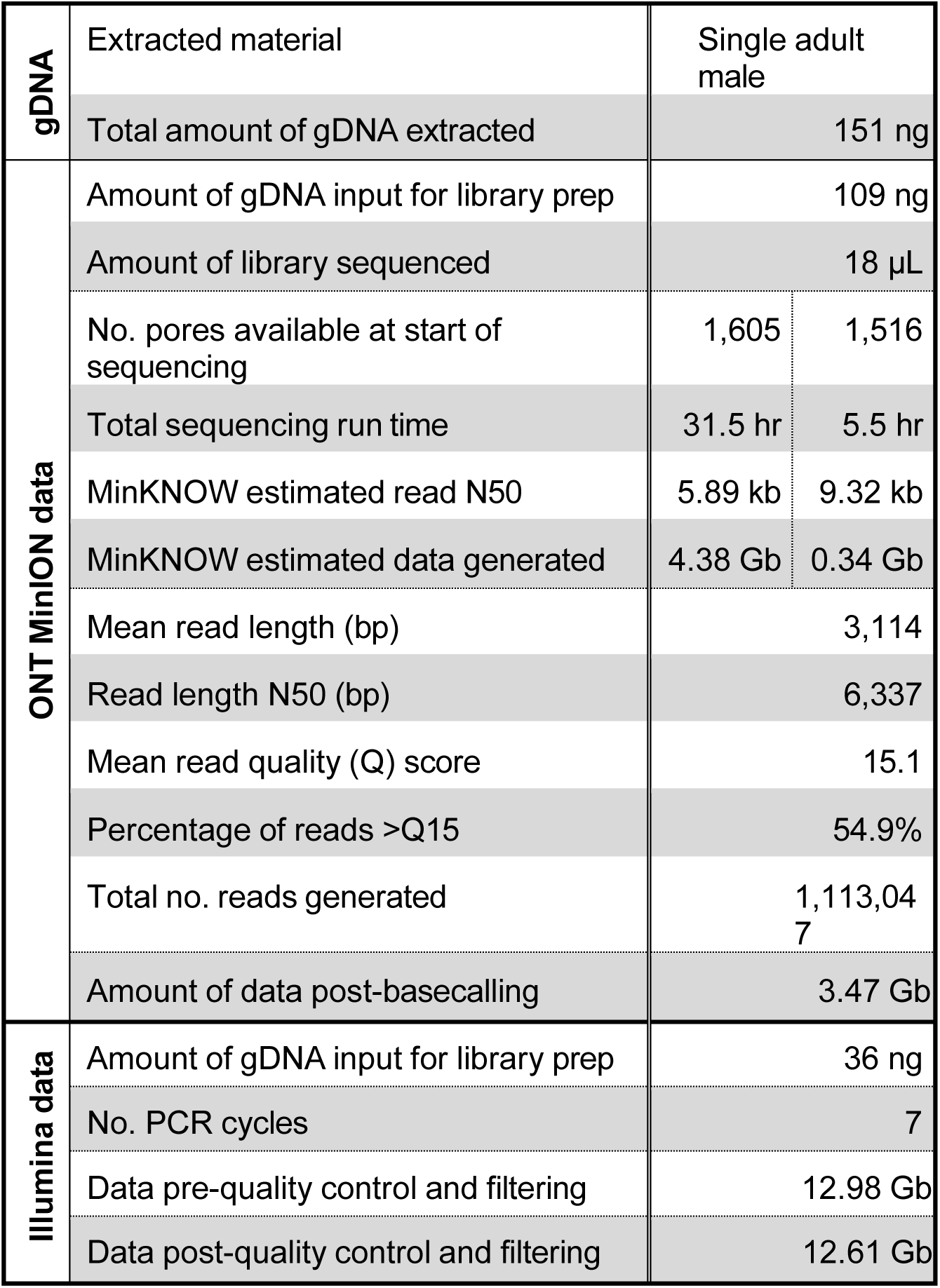
Nucleic acid extraction and Oxford Nanopore Technologies (ONT) MinION and Illumina next-generation sequence data generation for *Necator americanus*. Split cells indicate values for first (left) versus second (right) ONT MinION flow cells utilized.

|  |  |  |  |
| --- | --- | --- | --- |
| **gDNA** | Extracted material | Single adult male |  |
|  | Total amount of gDNA extracted | 151 ng |  |
| **ONT MinION data** | Amount of gDNA input for library prep | 109 ng |  |
|  | Amount of library sequenced | 18 µL |  |
|  | No. pores available at start of sequencing | 1,605 | 1,516 |
|  | Total sequencing run time | 31.5 hr | 5.5 hr |
|  | MinKNOW estimated read N50 | 5.89 kb | 9.32 kb |
|  | MinKNOW estimated data generated | 4.38 Gb | 0.34 Gb |
|  | Mean read length (bp) | 3,114 |  |
|  | Read length N50 (bp) | 6,337 |  |
|  | Mean read quality (Q) score | 15.1 |  |
|  | Percentage of reads >Q15 | 54.9% |  |
|  | Total no. reads generated | 1,113,047 |  |
|  | Amount of data post-basecalling | 3.47 Gb |  |
| **Illumina data** | Amount of gDNA input for library prep | 36 ng |  |
|  | No. PCR cycles | 7 |  |
|  | Data pre-quality control and filtering | 12.98 Gb |  |
|  | Data post-quality control and filtering | 12.61 Gb |  |

**S6 Table.** Top Mash hits for quality controlled Oxford Nanopore Technologies MinION and Illumina read datasets generated for *Necator americanus*.

|  | identity | shared-hashes | median-multiplicity | p-value | query-ID | query-comment |
| --- | --- | --- | --- | --- | --- | --- |
| Illumina | 0.986519 | 752/1000 | 37 | 0 | GCF_000507365.1_N_americanus_v1_genomic.fna.gz | [11864 seqs] NW_013562479.1 *Necator americanus* unplaced genomic scaffold N_americanus-1.0_Cont1, whole genome shotgun sequence [...] |
|  | 0.939885 | 272/1000 | 1 | 0 | GCF_000018665.1_ASM1866v1_genomic.fna.gz | NC_010002.1 *Delftia acidovorans* SPH-1, complete genome |
|  | 0.91333 | 149/1000 | 2 | 0 | GCF_000287295.1_ASM28729v1_genomic.fna.gz | NC_018417.1 *Candidatus Carsonella ruddii* HT isolate Thao2000, complete genome |
|  | 0.897826 | 104/1000 | 1 | 6.10E-229 | GCF_001320655.1_ASM132065v1_genomic.fna.gz | [147 seqs] NZ_BCBK01000001.1 *Pseudomonas* sp. NBRC 111134 DNA, contig: PPU21_CON00110, whole genome shotgun sequence [...] |
|  | 0.892599 | 92/1000 | 1 | 1.33E-197 | GCF_000090965.1_ASM9096v1_genomic.fna.gz | [2 seqs] NC_008513.1 *Buchnera aphidicola* BCc, complete genome [...] |
|  | 0.890433 | 57/652 | 64 | 8.47E-122 | GCF_000910735.1_ViralProj214366_genomic.fna.gz | NC_021929.1 *Malvastrum* leaf curl Philippines betasatellite, complete sequence |
|  | 0.889737 | 86/1000 | 2 | 3.30E-182 | GCF_001602025.1_ASM160202v1_genomic.fna.gz | [833 seqs] NC_031481.1 *Plasmodium gaboni* strain SY75 chromosome 1, whole genome shotgun sequence [...] |
|  | 0.887203 | 81/1000 | 2 | 1.57E-169 | GCF_000829155.1_ASM82915v1_genomic.fna.gz | NZ_AP013293.1 *Candidatus Sulcia muelleri* PSPU DNA, complete genome |
|  | 0.881654 | 71/1000 | 1 | 1.29E-144 | GCF_001578535.1_ASM157853v1_genomic.fna.gz | [28 seqs] NZ_LTBM01000001.1 *Candidatus Phytoplasma oryzae* isolate Mbita1 Pmin.contig.0_1, whole genome shotgun sequence [...] |
|  | 0.879223 | 67/1000 | 15 | 7.88E-135 | GCF_000871765.1_ViralProj19833_genomic.fna.gz | NC_009562.1 *Alternanthera* yellow vein virus satellite DNA beta, complete genome |
| ONT MinION | 0.986582 | 753/1000 | 9 | 0 | GCF_000507365.1_N_americanus_v1_genomic.fna.gz | [11864 seqs] NW_013562479.1 *Necator americanus* unplaced genomic scaffold N_americanus-1.0_Cont1, whole genome shotgun sequence [...] |
|  | 0.926662 | 202/1000 | 1 | 0 | GCF_000018665.1_ASM1866v1_genomic.fna.gz | NC_010002.1 *Delftia acidovorans* SPH-1, complete genome |
|  | 0.903605 | 119/1000 | 1 | 4.23E-280 | GCF_001320655.1_ASM132065v1_genomic.fna.gz | [147 seqs] NZ_BCBK01000001.1 *Pseudomonas* sp. NBRC 111134 DNA, contig: PPU21_CON00110, whole genome shotgun sequence [...] |
|  | 0.890433 | 57/652 | 25 | 4.34E-127 | GCF_000910735.1_ViralProj214366_genomic.fna.gz | NC_021929.1 *Malvastrum* leaf curl Philippines betasatellite, complete sequence |
|  | 0.889737 | 86/1000 | 5 | 3.45E-190 | GCF_000287295.1_ASM28729v1_genomic.fna.gz | NC_018417.1 *Candidatus Carsonella ruddii* HT isolate Thao2000, complete genome |
|  | 0.873204 | 58/1000 | 2 | 1.34E-118 | GCF_000090965.1_ASM9096v1_genomic.fna.gz | [2 seqs] NC_008513.1 *Buchnera aphidicola* BCc, complete genome [...] |
|  | 0.873204 | 58/1000 | 1 | 1.34E-118 | GCF_000223885.1_ASM22388v1_genomic.fna.gz | NC_015947.1 *Stenotrophomonas maltophilia* JV3, complete genome |
|  | 0.869907 | 36/672 | 30 | 5.39E-73 | GCF_000899295.1_ViralProj176433_genomic.fna.gz | NC_018671.1 *Sauropus* leaf curl disease associated DNA beta, complete genome |
|  | 0.868675 | 52/1000 | 2 | 5.48E-104 | GCF_001578535.1_ASM157853v1_genomic.fna.gz | [28 seqs] NZ_LTBM01000001.1 *Candidatus Phytoplasma oryzae* isolate Mbita1 Pmin.contig.0_1, whole genome shotgun sequence [...] |
|  | 0.86622 | 49/1000 | 3 | 8.48E-97 | GCF_000872085.1_ViralProj20983_genomic.fna.gz | NC_009893.1 Tomato yellow dwarf disease associated satellite DNA beta-[Kochi] virus, complete genome |

## Notes

### Competing Interest Statement

The authors have declared no competing interest.

